# Larvicidal activity of *Trichoderma atroviride* (Hypocreales: Hypocreaceae) against *Aedes albopictus* (Diptera: Culicidae)

**DOI:** 10.1101/2025.01.21.634114

**Authors:** David T. Hayes, Patil Tawidian, Ethan Schubert, Qing Kang, Amare J. Sumpter, Kristin Michel

## Abstract

Larviciding is an important part of effective integrated mosquito management. However, growing resistance to chemical and bacterial-based insecticides requires biocontrol agents with novel modes of action. Entomopathogenic fungi are good candidates for larval control due to their capability to infect mosquito larvae and their production of larvicidal compounds. In this study, we isolated a strain of *Trichoderma atroviride* from *Aedes albopictus* larvae collected in Manhattan, KS, USA. We used a laboratory-based microcosm assay to expose L3 *Ae. albopictus* larvae to *T. atroviride* conidia and culture supernatant treatments. Larvae were monitored daily for survival and development to pupae and adults. In addition, adult survival was monitored for ten-days post pupation, and wing lengths were measured to assess mosquito size. Our results revealed that *T. atroviride* culture supernatant was a potent larvicide towards *Ae. albopictus*. However, conidia by themselves were not larvicidal, indicating the major mode of killing was through toxicity exerted by the culture supernatant. We further showed that larval exposure to *T. atroviride* supernatant delayed larval development to pupae. Sex-specific adult survival was not affected by larval exposure to *T. atroviride*. However, wing length of male and female mosquitoes were reduced, indicating a reduction in adult mosquito body size as compared to the control. Taken together, this study identifies the culture supernatant from a novel strain of *T. atroviride* as a potent larvicide of *Ae. albopictus*, potentially expanding our toolbox for biological control of mosquitoes.

## Introduction

*Aedes albopictus*, originating from southeast Asia, has rapidly expanded its geographic range to all inhabited continents, due its use of the global transportation network (Hawley et al., 1987; Hahn et al., 2016; Kraemer et al., 2019; Swan et al., 2022). This mosquito species is a competent vector of medically important arboviruses, including dengue, chikungunya, yellow fever, and Zika viruses. These arboviruses can inflict significant morbidity, mortality, and economic burden particularly to low and middle-income countries (Diagne et al., 2021; Ogunlade et al., 2021; Chilakam et al., 2023; Roiz et al., 2024) and lack sufficient prophylactic or therapeutic interventions to effectively reduce pathogen transmission (Troppens, 2024). For these reasons, the World Health Organization introduced the Global Arbovirus Initiative that includes mosquito control as a core pillar of this initiative (WHO, 2024). Mosquito control of *Aedes* species, including *Ae. albopictus*, targets both larval and adult stages (Rose, 2001; Floore, 2006; Roiz et al., 2018; Fouet and Kamdem, 2019). Indeed, simultaneous targeting of both stages is shown to be most effective to reduce *Ae. albopictus* populations in high-density areas were breeding sites cannot feasibly be removed (Wilke et al., 2021).

Mosquito larvicides include chemical and biological insecticides with different modes of action (McGregor and Connelly, 2020; Karunaratne and Surendran, 2022; Rodrigues Dos Santos et al., 2023). However, resistance to traditionally used larvicides has been reported for several mosquito species including *Ae. albopictus* (Dame et al., 1998; Vontas et al., 2012; Marcombe et al., 2014; Su et al., 2019; Hassan et al., 2021; Lopez et al., 2024). While resistance in mosquito populations towards the bioinsecticide *Bacillus thuringiensis* thus far remains limited (Silva-Filha et al., 2021), resistance to *Lysinibacillus sphaericus* has been reported in field collected *Culex* species (Su et al., 2018; Su et al., 2019). Thus, identifying novel biological control agents with different modes of action, such as entomopathogenic fungi (EPF) and their secreted metabolites, may offer new avenues for sustainable larval reduction.

EPF employ multiple methods to kill mosquitoes, such as actively infecting and propagating within the mosquito or through the secretion of entomotoxins (Shen et al., 2020). Fungi that kill mosquitoes by active colonization and metabolite secretion include Hypocrealean fungi such as *Beauveria bassiana* and *Metarhizium anisopliae*, both of which have shown potential for mosquito larvae control in laboratory and semi-field studies (Clark et al., 1967; Clark et al., 1968; Bukhari et al., 2010; Farida et al., 2018). In addition, several species in the Hypocrealean genus *Trichoderma* have shown mosquito larvicidal potential (Poveda, 2021). *Trichoderma* species are widely ubiquitous in soil and aquatic environments (Hu et al., 2020; Poveda, 2021) and are frequently used as biological control agents of numerous agriculturally relevant fungal phytopathogens (Asad, 2022). Several *Trichoderma* species, including *Trichoderma atroviride*, act as entomopathogens of insects through direct parasitism (Anwar et al., 2016; Ghosh and Pal, 2016; Rodríguez-González et al., 2016) or through the secretion of entomotoxic secondary metabolites (Govindarajan et al., 2005; Sundaravadivelan and Padmanabhan, 2014; Singh and Prakash, 2015; Podder and Ghosh, 2019).

In this study, we assessed the mosquito control potential of a *T. atroviride* strain, initially isolated from *Ae. albopictus* L4 larvae. Modifying a previously established laboratory-based experimental design (Tawidian et al., 2023), we conducted bioassays exposing L3 *Ae. albopictus* larvae to *T. atroviride*. Several life history parameters were recorded, including daily survival of larvae, pupae, and adults, development time, and adult size and survival following larval exposure. By using conidia and culture supernatants in separate treatments, we also assessed whether *T. atroviride* killing of *Ae. albopictus* larvae is due to entomopathogenicity or entomotoxicity.

## Methods

### Isolation and molecular identification of *T. atroviride* from *Ae. albopictus* larvae

*Trichoderma atroviride* was isolated from field collected L4 *Ae. albopictus* larvae as described previously (Tawidian et al., 2023). Briefly, L4 *Ae. albopictus* larvae were collected from one location (N 39° 11ʹ35.5ʺ, W 96° 34ʹ15.7ʺ) in Manhattan, KS from mosquito oviposition cups lined with seed germination paper. *Trichoderma atroviride* was isolated by washing individual larvae six times with 1% PBS and grinding them in 50 µL of sterile water. Each larval suspension was plated on a potato dextrose agar (PDA, Millipore, MA, USA) supplemented with 100 mg/L ampicillin and 25 mg/L chloramphenicol to prevent bacterial contamination. PDA plates were incubated at room temperature in the dark for 5 days. Unique fungal morphotypes were then isolated for fungal molecular identification and propagation.

To determine the identity of the isolated fungi, we extracted total genomic DNA from fungal mycelia and conidia using the DNeasy PowerSoil kit (MoBio Laboratory, CA, USA) according to the manufacturer’s instructions. We then amplified the fungal Internal Transcribed Spacer (ITS) marker using forward primer ITS1f (5’-CTTGGTCATTTAGAGGAAGTAA-3’) (Gardes and Bruns, 1993) and reverse primer ITS4 (5’-TCCTCCGCTTATTGATATGC-3’) (White et al., 1990) according to the protocol described previously (Ihrmark et al., 2012). PCR products were purified using a Qiaquick® PCR purification kit (Qiagen, Germany). The purified PCR amplicons were sequenced by Sanger sequencing (GeneWiz, NJ, USA), blasted to the NCBI nucleotide database and compared to similar sequences. The ITS region sequence was submitted to GenBank and assigned the accession #OP692519.1.

### Mosquito maintenance

*Aedes albopictus* (initially collected in Missouri and colonized by the Illinois Natural History Survey) mosquitoes were reared at 27°C and 80% relative humidity using a 12L:12D photoperiod cycle. To hatch *Ae. albopictus* eggs, egg papers were introduced to water containing 0.36 g/L of CM0001 Nutrient Broth (Zheng et al., 2015). Thereafter, larvae were fed daily on a suspension of baker’s yeast (Fleischman’s Active dry Yeast, 6.6 g/L) and fish food (Tetramin Tropical Flakes, 13.6 g/L). Pupae were then transferred to adult emergence cages. Adult mosquitoes were provided a sugar solution containing 8% fructose supplemented with 2.5 mM PABA *ad libitum*.

### Propagation of *T. atroviride* on wheat

Sterilized wheat kernels were used as a growth medium for *T. atroviride* to achieve the desired concentration and volume of conidia for exposure bioassays. Fungal inoculation jars were prepared by combining 100 g of dry wheat kernels (Kansas wheat variety ‘Everest’, USDA, Manhattan, KS, USA) with 40 mL of Milli-Q water in 32 oz glass mason jars. The mason jars were then covered with four layers of cheesecloth, two layers of paper towel, and a layer of aluminum foil. The prepared mason jars were sterilized by autoclaving at 121°C and 15 psi for 40 minutes and left to dry overnight before inoculation with *T. atroviride* conidia. *T. atroviride* conidia were harvested from 14-day old PDA plates and diluted to 10^7^ conidia/mL using sterile Milli-Q water. 10 mL of 10^7^ conidia/mL suspension were added to the sterile wheat kernels through puncturing the paper towel and cheesecloth layers with a sterile syringe under aseptic conditions. Wheat kernels were agitated by shaking and the jars were incubated in the dark at room temperature for 14 days.

### Harvesting *T. atroviride* conidia from wheat kernels

Conidia were harvested by transferring wheat kernels in batches to sterile 100 mL glass bottles (Fisher Scientific, NH, USA) containing sterile Milli-Q water. Wheat kernels were then shaken manually for 10-15 seconds to detach the conidia from the kernels. The suspension was then filtered through two layers of cheesecloth into a sterile 50 mL Falcon^®^ tube (Fisher Scientific, NH, USA) to separate conidia from mycelia and wheat kernels. The subsequent conidial suspension was quantified using a hemocytometer and adjusted to 10^6^, 10^7^, and 10^8^ conidia/mL concentrations, respectively. The 10^8^ conidia/mL suspension was used to prepare two additional treatments of culture supernatant (SUP) and autoclaved culture supernatant (AC SUP). To prepare the culture supernatant, the 10^8^ conidia/mL suspension was centrifuged at 2,500 g for 20 minutes and the supernatant was collected by decanting, reducing its conidial concentration to approximately 4,000 conidia/mL. Some of the resulting supernatant treatment was then autoclaved at 121°C and 15 psi for 40 mins to ensure the killing of all remaining conidia and inactivating any heat-labile compounds. Lastly, a treatment consisting of fungal conidia in the absence of fungal supernatant (referred to as washed conidia) was prepared by washing the conidial pellet, leftover from the supernatant preparation, with sterile milli-Q water. The suspension was vortexed well to create a homogenous suspension, centrifuged at 2,500 g for 20 minutes, and the conidial pellet was resuspended in 20-30 mL sterile milli-Q water. The washing steps were repeated two additional times, and the final washed conidial suspension was adjusted to 10^8^ conidia/mL. The 10^8^ conidia/mL and washed conidia treatments were evaluated for germination prior to exposure by plating 100 µl of each suspension onto PDA. Both treatments had a mean germination between 94-96% prior to larval exposure.

### Exposure bioassay

To assess the impact of *Ae. albopictus* larval exposure to *T. atroviride* on mosquito survival and development, we performed exposure bioassays conducted in 24-well plates using six fungal treatments and a water-only control (Fig. S1). Individual microcosms contained one L3 larva and 1 mL of fungal or control treatments, including 10^6^, 10^7^, and 10^8^ conidia, supernatant and autoclaved supernatant, and washed 10^8^ conidia, or a water-only control. Larvae were fed daily by adding 20 µl of food slurry (6.6 g/L baker’s yeast, 13.6 g/L fish food in Milli-Q water) to each well. Larvae were monitored daily for death, survival, and pupation. Pupae were collected daily and separated based on pupation date and treatment. Pupae were washed in water and placed into emergence cups. Emergence cups were monitored daily for pupal survival, death, and eclosion of male and female adults. Adults were fed using cotton balls soaked in a fructose mixture (8% fructose with 2.5 mM PABA in Milli-Q water) placed on top of the emergence cups. Male and female mosquitoes were monitored daily for death up to 10 days post pupation.

To assess the impact of *Ae. albopictus* L3 larval exposure to *T. atroviride* on mosquito size, male and female adults from each treatment were killed by placing the emergence cups at *-*20°C overnight. A single wing from each individual mosquito was removed and its length determined as described previously (Romoli and Gendrin, 2020), by imaging using 32x magnification (Leica MZ7.5, Leica Microsystems, Germany) and measuring pixels using Image J (version 1.53t; Java 1.8.0_345 [64-bit], Schneider et al., 2012), estimating the distance between the alula notch to the intersection of the radium 3 vein and the outer margin, excluding fringe scales.

The first set of bioassays, hereby referred to as experiment 1, evaluated the efficacy of *T. atroviride* conidia and supernatant towards *Ae. albopictus* larvae by including the following treatments 10^6^, 10^7^, and 10^8^ conidia/mL, supernatant and autoclaved supernatant from the 10^8^ conidia/mL treatment, and a water-only control. A separate experiment, hereafter referred to as experiment 2, evaluated larvicidal activity of *T. atroviride* conidia in the absence of fungal supernatant by exposing larvae to the washed conidia treatment in parallel with the 10^8^ conidia/mL treatment, supernatant and autoclaved supernatant from the 10^8^ conidia/mL treatment, and water-only control. For both experiments, each treatment was performed in three 24-well plates (n = 72 larvae). Experiments 1 and 2 were repeated 3 and 4 times, respectively, using different batches of mosquitoes and fungal cultures.

### Conidial concentration and viability within larval microcosms

To determine the daily fluctuation in the viability and concentration of conidia within in the larval microcosm during entire exposure time, we quantified conidia and their germination percentages daily for the 10^8^ conidia/mL and washed conidia treatments of experiment 2. Each day, we collected eight pools of conidial suspensions/treatment (each pool containing 20 µL from six individual microcosms). For each pool, conidia were counted using a hemocytometer. To determine conidial viability in each pool, germination percentages were assessed on PDA plates supplemented with 100 mg/L ampicillin and 25 mg/L chloramphenicol. Germination percentages were evaluated 18 hours post-inoculation, by counting a total of 300 conidia per pool under optical microscopy (400x). Conidia were considered germinated if the length of its germ tube was at least twice the length of the diameter of the conidium (Francisco et al., 2006). Conidia germination did not significantly decrease throughout the course of the exposure and was greater than 95% for the duration of larval exposure (Fig. S2).

### Qualitative assessment of conidia ingestion

To qualitatively assess if conidia were ingested, we exposed L3 larvae to 10^8^/mL acridine orange (AO) labelled conidia. Washed conidia at a 10^8^ conidia/mL concentration were resuspended in 200 μg/mL AO and incubated for 1 h at room temperature. Conidia were washed in sterile milli-Q water three times as described above, and resuspended in sterile milli-Q water at a final concentration of 10^8^ conidia/mL. Larvae were exposed to the AO-labelled conidia in microcosms and images were taken 24 h post exposure using a fluorescent scope (Leica EL6000, North Central Instruments, MO, USA) fitted with an eGFP filter.

### Fungal sporulation from larval carcasses

To determine whether *T. atroviride* can use larval carcasses as a growth substrate, killed larvae were placed on water agar and observed for fungal sporulation. Dead larvae from the 10^8^ conidia/mL and washed 10^8^ conidia/mL treatments were surface sterilized in 10% bleach for 2 minutes and rinsed twice in sterile Milli-Q water for 2 minutes under aseptic conditions. Each surface-sterilized larva was then placed into an individual well containing 1mL of water agar (15 g agar/L milli-Q water, supplemented with 100 mg/L ampicillin and 25 mg/L chloramphenicol to prevent bacterial growth) using 24-well plates. Inoculated plates were incubated at room temperature in the dark and were monitored for *T. atroviride* sporulation from dead larvae. Brightfield images were taken 13 days post larval death using a second-generation iPhone SE.

### Statistical analyses

Most statistical analyses were executed using the MIXED, LOGISTIC, PHREG and LIFETEST procedures in SAS/STAT® software, version 9.4 (SAS Institute Inc., NC, USA) and were conducted as previously described (Tawidian et al., 2023). In brief, we assessed the L1 to pupae and L1 to sex-specific adult survival using conditional logistic regression model with fungal treatment as fixed effect and trial forming strata (dataset found in Table S1). We used a linear mixed model with fungal treatment and trial as fixed effects to assess larval developmental time as measured by time to pupation and time to eclosion. Least squares means (LSMs) and standard errors (SEMs) were reported for each treatment. Adult survival post-eclosion was analyzed using Cox’s proportional hazard model with fungal treatment as the fixed effect and trial forming strata. Some treatments were excluded from analysis due to the lack of larvae developing to adults or the absence of adult death. Survival curves were obtained using Kaplan-Meier estimator.

Linear regression was performed to determine if the conidia concentration and germination percentage of the washed conidia treatments decreased throughout the course of the fungal exposure. Adult male and female wing lengths were analyzed using one-way ANOVA with Tukey’s multiple comparisons test for experiment 1 and t-test for experiment 2 on GraphPad prism (version 10.1.1, GraphPad Software, MA, USA). Adult sex ratio and pupal mortality were assessed using Fisher’s exact test comparing fungal treatments to water-only control using GraphPad Prism.

## Results

### Morphological and molecular characterization of the *T. atroviride* isolate MHK

A fungal strain was isolated from *Ae. albopictus* L4 mosquito larvae collected from human-made containers in Manhattan, KS, USA. Colony morphology on PDA varied across time. Early colonies (3 days post-inoculation) had white cottony filamentous growth on the surface and on the reverse side. Older colonies (7 days post-inoculation) developed yellow and light and dark green pigments on the surface, typical for *Trichoderma* spp., while on the reverse side the growth remained white (Fig. 1A and B). Conidiophores were repeatedly and irregularly branched, with clusters of flask-shaped phialides with terminal conidia. Conidia were 2.5-3.3 µm in size, hyaline, and globose to subglobose in shape, characteristic of *Trichoderma* spp. (Steyaert et al., 2010, Fig. 1C). Using sequencing-based approaches, we characterized *Trichoderma* to the species level. Nucleotide sequence comparison to the nonredundant GenBank database revealed 99.8% sequence identity to a known *Trichoderma atroviride* isolate.

**Fig. 1.**
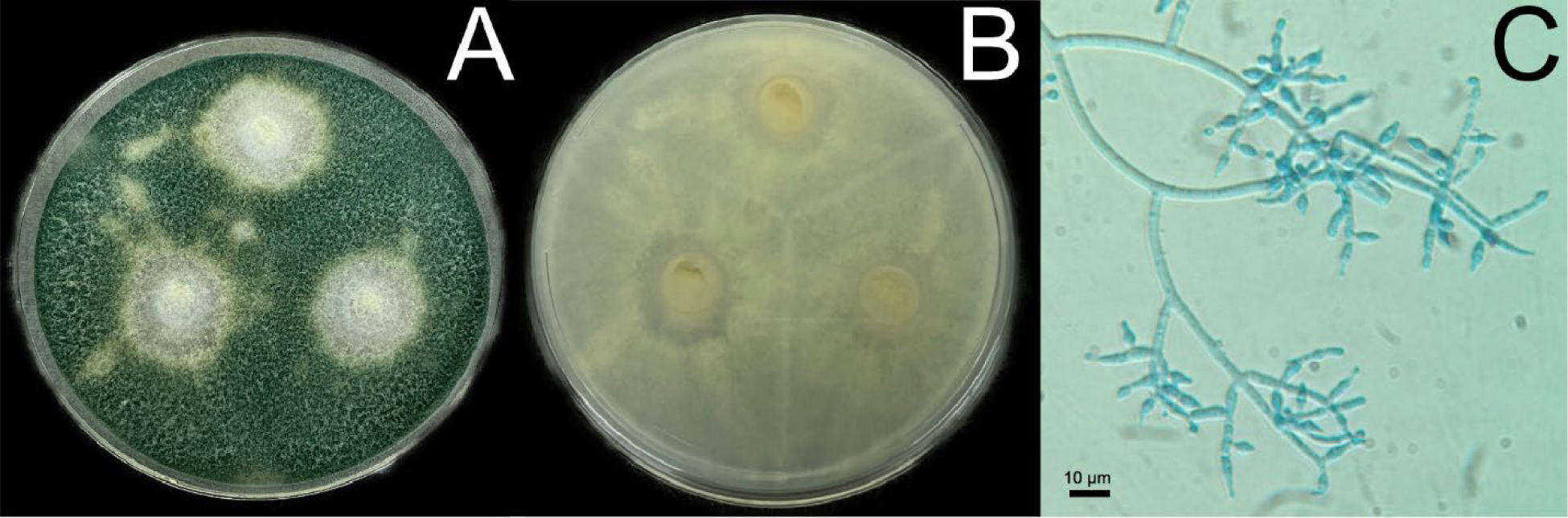
Colony and microscopic morphology of *T. atroviride*. **(A)** Fungal growth on the surface and **(B)** reverse side of *T. atroviride* on PDA. **(C)** Microscopic morphology of *T. atroviride* hyphae, conidia, conidiophores, and phialides. Scale bar, 10 μm.

### Trichoderma atroviride conidia are consumed by Ae. albopictus larvae

To assess qualitatively conidia consumption by *Ae. albopictus* larvae, we exposed L3 larvae to AO-labelled conidia. We found AO fluorescence in larval guts up to the fourth abdominal segment 24 h after exposure, demonstrating that *Ae. albopictus* larvae did consume *T. atroviride* conidia in our experimental set-up (Fig. S3). We next tested whether larval consumption reduced the total number of conidia in their environment. To do so, we quantified the conidial concentrations in the microcosms daily over the exposure period. Conidial concentrations in the washed conidial and 10^8^ conidia/mL treatments decreased from 10^8^ conidia/mL to 4.8x10^7^ and 6.3x10^7^ conidia/mL after 24 h of exposure, respectively (Fig. S4). Washed conidial counts decreased further over time, reaching 3.3x10^7^ conidia/mL 4 days post exposure (Fig. S4A). In the 10^8^ conidia/mL treatment too many larvae had died on day 1 to evaluate conidia concentration on subsequent days.

### *T. atroviride* supernatant but not conidia killed *Ae. albopictus* larvae

To assess the larvicidal impact of *T. atroviride* on *Ae. albopictus* larvae, survival proportions and odds ratios of L1 to pupa survival were compared across fungal treatments and the water-only control (Table 1, Fig. 2A and C). In experiments 1 and 2, we observed a significant reduction in larval survival to the pupal stage when exposed to the 10^8^ conidia/mL treatment. Only 25.0% and 2.4% of larvae exposed to this treatment reached the pupal stage in experiments 1 and 2, respectively, as compared to the water-only control, in which 96.8% and 94.4% survived. In both experiments, larvae had a significantly reduced odds ratio (OR) when exposed to a high dose of *T. atroviride* conidia compared to water (OR = 0.01, *P* < 0.001). The significant reduction in larval survival was dose dependent. Exposure to lower conidia concentrations did not reduce larval survival; 96.8% and 94.4% of larvae pupated when exposed to 10^6^ and 10^7^ conidia/mL treatments, respectively. The 10^6^ and 10^7^ conidia/mL treatments were therefore excluded from experiment 2.

**Fig. 2.**
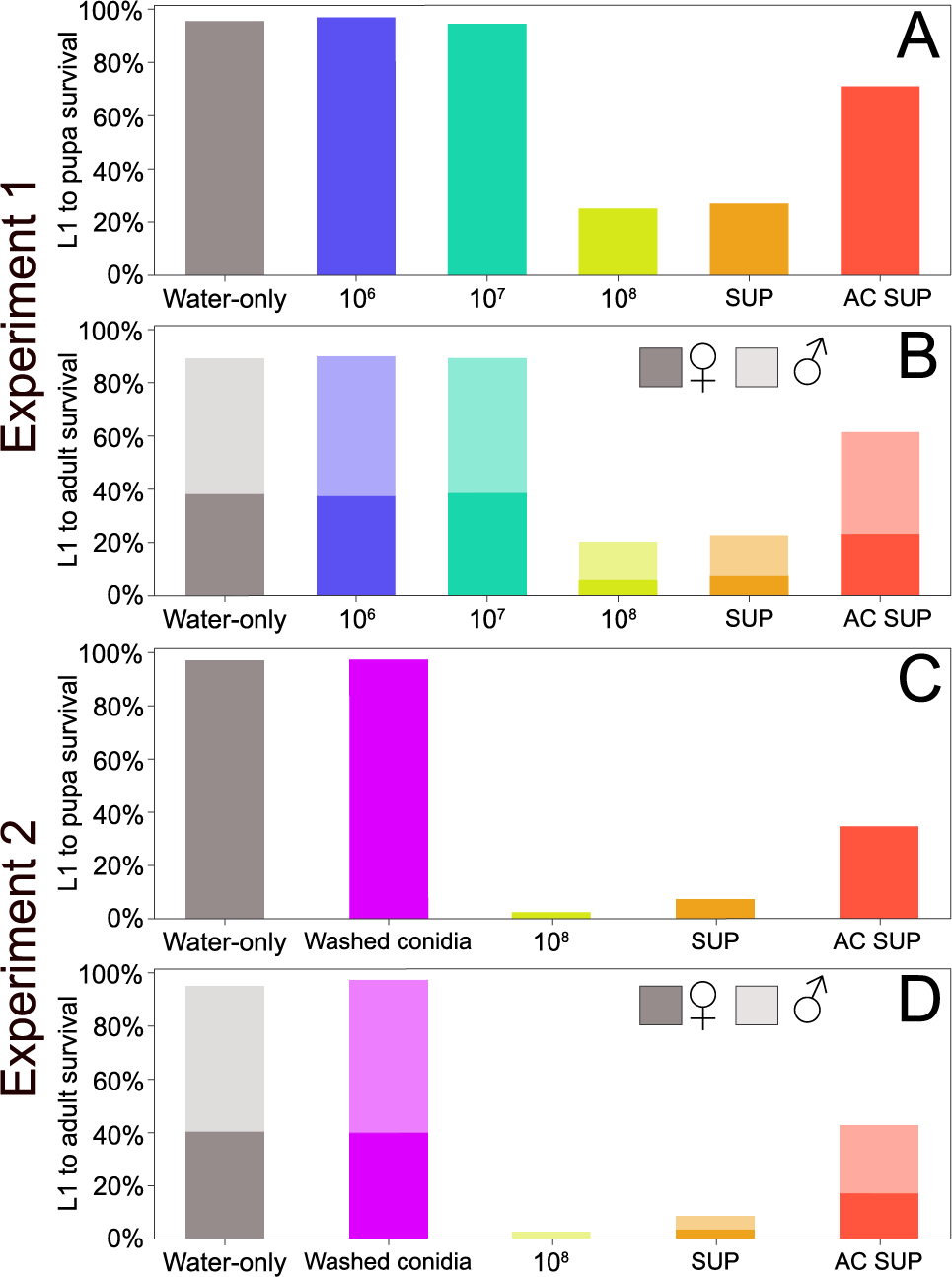
L1 to pupa and L1 to sex-specific adult survival percentages of *Ae. albopictus* larvae exposed to *T. atroviride*. (A and C) L1 to pupa survival and (**B and D)** L1 to adult survival of *Ae. albopictus* larvae following exposure to *T. atroviride* in experiments 1 and 2, respectively. Darker bars indicate survival to adult female, while lighter bars represent survival to adult male.

**Table 1.**
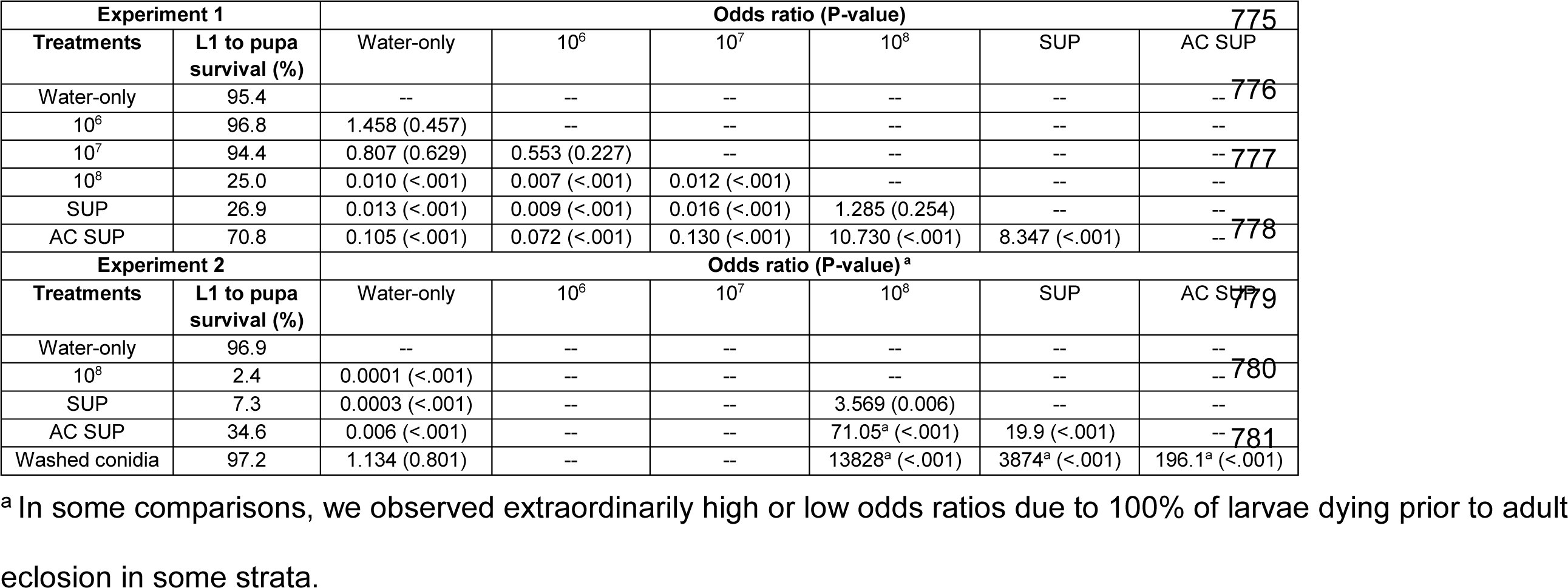
Odds ratios of *Ae. albopictus* L1 to pupa survival following exposure to *T. atroviride*.

To determine if *T. atroviride* culture supernatant contributes to larvicidal activity, we exposed *Ae. albopictus* larvae to supernatant and autoclaved supernatant treatments (Table 1, Fig. 2A and C). Similar to the 10^8^ conidia/mL treatment, the *T. atroviride* supernatant significantly reduced larval survival compared to the water-only control (OR = 0.013, *P <* 0.001), indicating that entomotoxins present in the culture supernatant contributed to larvicidal activity. We then autoclaved the culture supernatant to inactivate heat-labile molecules and kill any remaining conidia. Exposure to the autoclaved supernatant treatment had an increased larval survival compared to the supernatant treatment (OR = 8.347, *P <* 0.001 for experiment 1 and OR = 19.9, *P <* 0.001 for experiment 2). However, autoclaving did not completely inactivate the larvicidal activity of *T. atroviride* supernatant (OR = 0.105, *P <* 0.001 for experiment 1 and OR = 0.006, *P <* 0.001 for experiment 2 compared to the water-only control).

We then set out to determine if *T. atroviride* conidia by themselves had any larvicidal activity (Table 1, Fig. 2C). To do so, we exposed *Ae. albopictus* larvae to a washed conidia treatment containing 10^8^ conidia/mL, but in the absence of fungal supernatant. We observed no impact to *Ae. albopictus* survival as 97.2% of larvae exposed to the washed conidia treatment reached the pupal stage (OR = 1.134, *P* = 0.801 compared to water-only). Thus, *T. atroviride* conidia lack larvicidal ability.

Although, *T. atroviride* conidia in the absence of supernatant had no larvicidal activity, the conidia may use the larval carcass as a growth substrate for sporulation. To evaluate this, the carcasses of larvae that died following exposure to the 10^8^ conidia/mL and washed conidia treatments were surface sterilized and plated onto agar lacking any nutrients. We observed white, cottony filamentous growth protruding out and surrounding the larval carcass with green pigmentation at the growth edge (Fig. S5), demonstrating *T. atroviride* growth and sporulation.

To determine whether *T. atroviride* also reduced pupal survival when initially exposed to L3 larvae, we plotted L1 to adult survival and compared the percentage of L1 to pupa survival to that of L1 to adult survival (Table S2, Fig 2). In water-only treatments, 6.4% and 3.6% of mosquitoes died at the pupal stage in experiments 1 and 2, respectively. Similar pupal mortality was observed when larvae were exposed to the 10^6^, 10^7^, and washed conidia treatments. The few larvae that survived and pupated after exposure to supernatant and autoclaved supernatant, were twice as likely to die in the pupal stage compared to the water-only control (Table S2).

### *Trichoderma atroviride* differentially impacted sex-specific immature survival

To evaluate if *T. atroviride* conidia and supernatant treatments had a sex-specific killing effect, we individually plotted L1 to adult survival proportions for female and male adults (Table 2, Fig. 2B and D) and directly compared the adult sex ratios between treatments (Table 3). Following exposure to the water-only control, 51.3-55% of *Ae. albopictus* larvae eclosed to adult males and 37.9-40.1% to adult females. We observed a reduction in survival to both adult males and females when exposed to the 10^8^ conidia/mL, supernatant, or autoclaved supernatant treatments (Table 2). For example, in experiment 1, larval exposure to the 10^8^ conidia/mL treatment resulted in only 14.8% of larvae surviving to adult males (OR = 0.146, *P <* 0.001) and 5.6% (OR = 0.081, *P <* 0.001) surviving to adult females.

**Table 2.**
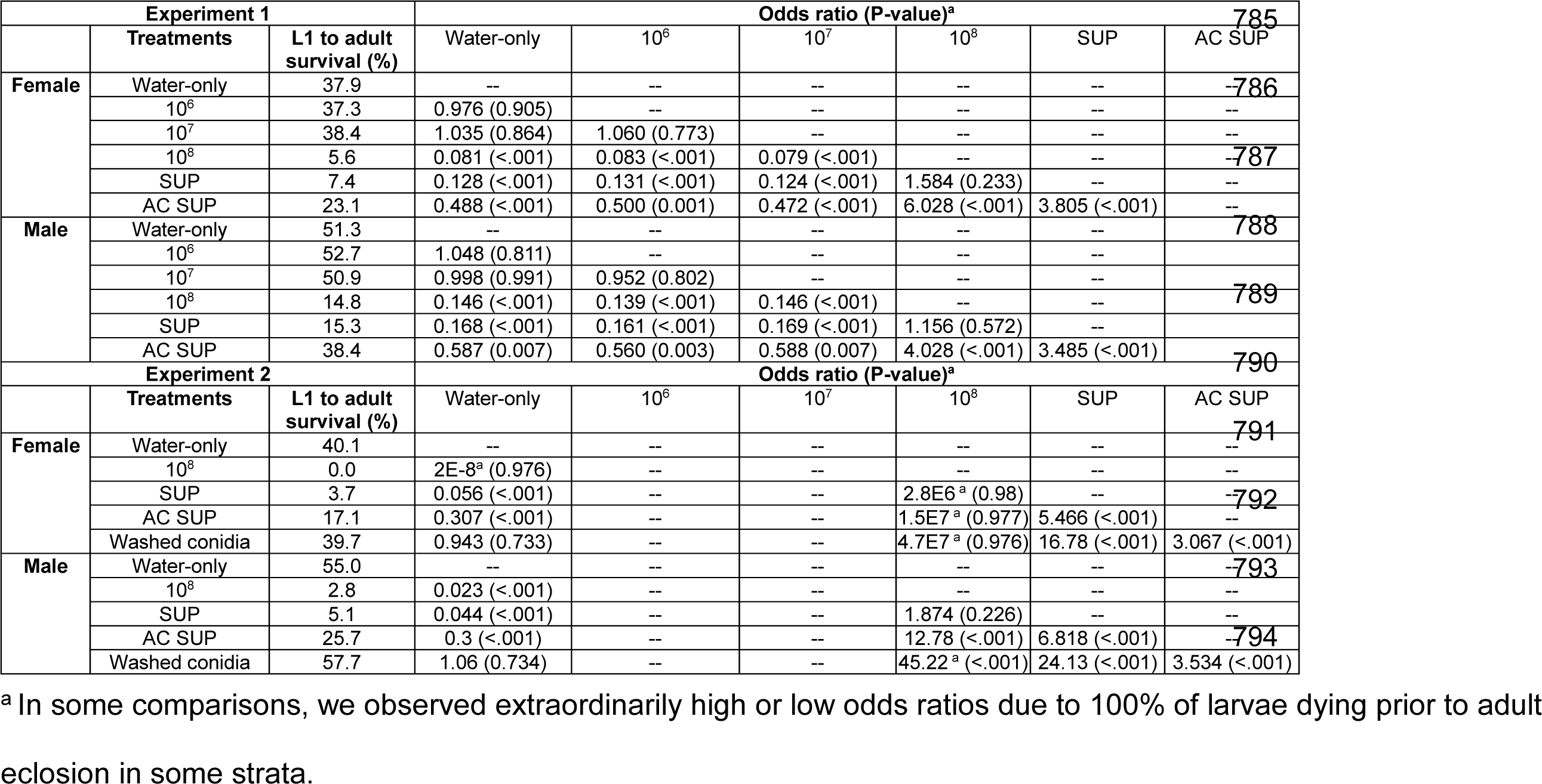
Odds ratios of *Ae. albopictus* L1 to adult survival following exposure to *T. atroviride*.

| Experiment 1 |  |  | Odds ratio (P-value)<sup>a</sup> |  |  |  |  |  | 785 |
| --- | --- | --- | --- | --- | --- | --- | --- | --- | --- |
|  | Treatments | L1 to adult survival (%) | Water-only | 10<sup>6</sup> | 10<sup>7</sup> | 10<sup>8</sup> | SUP | AC SUP |  |
| Female | Water-only | 37.9 | -- | -- | -- | -- | -- | -- | -786 |
|  | 10<sup>6</sup> | 37.3 | 0.976 (0.905) | -- | -- | -- | -- | -- |  |
|  | 10<sup>7</sup> | 38.4 | 1.035 (0.864) | 1.060 (0.773) | -- | -- | -- | -- |  |
|  | 10<sup>8</sup> | 5.6 | 0.081 (<.001) | 0.083 (<.001) | 0.079 (<.001) | -- | -- | -- | -787 |
|  | SUP | 7.4 | 0.128 (<.001) | 0.131 (<.001) | 0.124 (<.001) | 1.584 (0.233) | -- | -- |  |
|  | AC SUP | 23.1 | 0.488 (<.001) | 0.500 (0.001) | 0.472 (<.001) | 6.028 (<.001) | 3.805 (<.001) | -- |  |
| Male | Water-only | 51.3 | -- | -- | -- | -- | -- | -- | 788 |
|  | 10<sup>6</sup> | 52.7 | 1.048 (0.811) | -- | -- | -- | -- | -- |  |
|  | 10<sup>7</sup> | 50.9 | 0.998 (0.991) | 0.952 (0.802) | -- | -- | -- | -- |  |
|  | 10<sup>8</sup> | 14.8 | 0.146 (<.001) | 0.139 (<.001) | 0.146 (<.001) | -- | -- | -- | 789 |
|  | SUP | 15.3 | 0.168 (<.001) | 0.161 (<.001) | 0.169 (<.001) | 1.156 (0.572) | -- | -- |  |
|  | AC SUP | 38.4 | 0.587 (0.007) | 0.560 (0.003) | 0.588 (0.007) | 4.028 (<.001) | 3.485 (<.001) | -- | 790 |
| Experiment 2 |  |  | Odds ratio (P-value)<sup>a</sup> |  |  |  |  |  |  |
|  | Treatments | L1 to adult survival (%) | Water-only | 10<sup>6</sup> | 10<sup>7</sup> | 10<sup>8</sup> | SUP | AC SUP |  |
| Female | Water-only | 40.1 | -- | -- | -- | -- | -- | -- | 791 |
|  | 10<sup>8</sup> | 0.0 | 2E-8<sup>a</sup> (0.976) | -- | -- | -- | -- | -- |  |
|  | SUP | 3.7 | 0.056 (<.001) | -- | -- | 2.8E6<sup>a</sup> (0.98) | -- | -- | -792 |
|  | AC SUP | 17.1 | 0.307 (<.001) | -- | -- | 1.5E7<sup>a</sup> (0.977) | 5.466 (<.001) | -- |  |
|  | Washed conidia | 39.7 | 0.943 (0.733) | -- | -- | 4.7E7<sup>a</sup> (0.976) | 16.78 (<.001) | 3.067 (<.001) |  |
| Male | Water-only | 55.0 | -- | -- | -- | -- | -- | -- | -793 |
|  | 10<sup>8</sup> | 2.8 | 0.023 (<.001) | -- | -- | -- | -- | -- |  |
|  | SUP | 5.1 | 0.044 (<.001) | -- | -- | 1.874 (0.226) | -- | -- |  |
|  | AC SUP | 25.7 | 0.3 (<.001) | -- | -- | 12.78 (<.001) | 6.818 (<.001) | -- | -794 |
|  | Washed conidia | 57.7 | 1.06 (0.734) | -- | -- | 45.22<sup>a</sup> (<.001) | 24.13 (<.001) | 3.534 (<.001) |  |

<sup>a</sup> In some comparisons, we observed extraordinarily high or low odds ratios due to 100% of larvae dying prior to adult eclosion in some strata.

**Table 3.** Sex ratio of *Ae. albopictus* adults that survived larval fungal exposure.

| Experiment 1 | Treatment | Males:Females | Ratio | P |
| --- | --- | --- | --- | --- |
|  | Water-only | 110:81 | 1.36:1 | -- |
|  | 10<sup>6</sup> | 110:78 | 1.41:1 | 0.917 |
|  | 10<sup>7</sup> | 109:82 | 1.33:1 | >0.999 |
|  | 10<sup>8</sup> | 40:13 | 3.08:1 | 0.025 |
|  | SUP | 33:16 | 2.06:1 | 0.255 |
|  | AC SUP | 83:50 | 1.66:1 | 0.421 |
| Experiment 2 | Treatment | Males:Females | Ratio | P |
|  | Water-only | 154:111 | 1.39:1 | -- |
|  | 10<sup>8</sup> | 6:0 | -- | -- |
|  | SUP | 11:8 | 1.38:1 | >0.999 |
|  | AC SUP | 55:38 | 1.45:1 | 0.903 |
|  | Washed conidia | 151:117 | 1.29:1 | 0.726 |

To determine whether the reduction in survival to eclosion affected females or males disproportionately, we evaluated adult sex ratios across treatments (Table 3). Larvae exposed to the water-only control developed to adults at a 1.36:1 male:female ratio. This ratio was similar across the two lower conidia dose treatments (1.41:1 and 1.33:1 ratio for the 10^6^ and 10^7^ conidia/mL). However, larvae that survived to adulthood following exposure to the 10^8^ conidia/mL treatment were significantly male-skewed as 3 males were observed for every 1 female (*P* = 0.025 compared to water-only control).

### Trichoderma atroviride delayed Ae. albopictus larval development

Time to pupation and time to eclosion parameters were used to determine the impact of *T. atroviride* on *Ae. albopictus* development (Table 4, Fig. 3). When exposed to the water-only control, *Ae. albopictus* larvae developed to pupae in 6.4 days and adults eclosed 2.2 days later. Larval development time was not delayed for larvae exposed to the two lowest conidial doses compared to the water-only control. Larval development to pupae was at 6.5 days and adults eclosed 2.2-2.3 days later for the 10^6^ and 10^7^ conidia/mL doses, respectively (Table 4, Fig. 3A and B). Similarly, the washed conidia treatment did not significantly delay larval development (0.2 day delay, *P* = 0.296) nor time to adult eclosion (Table 4, Fig. 3C and D).

**Fig. 3.**
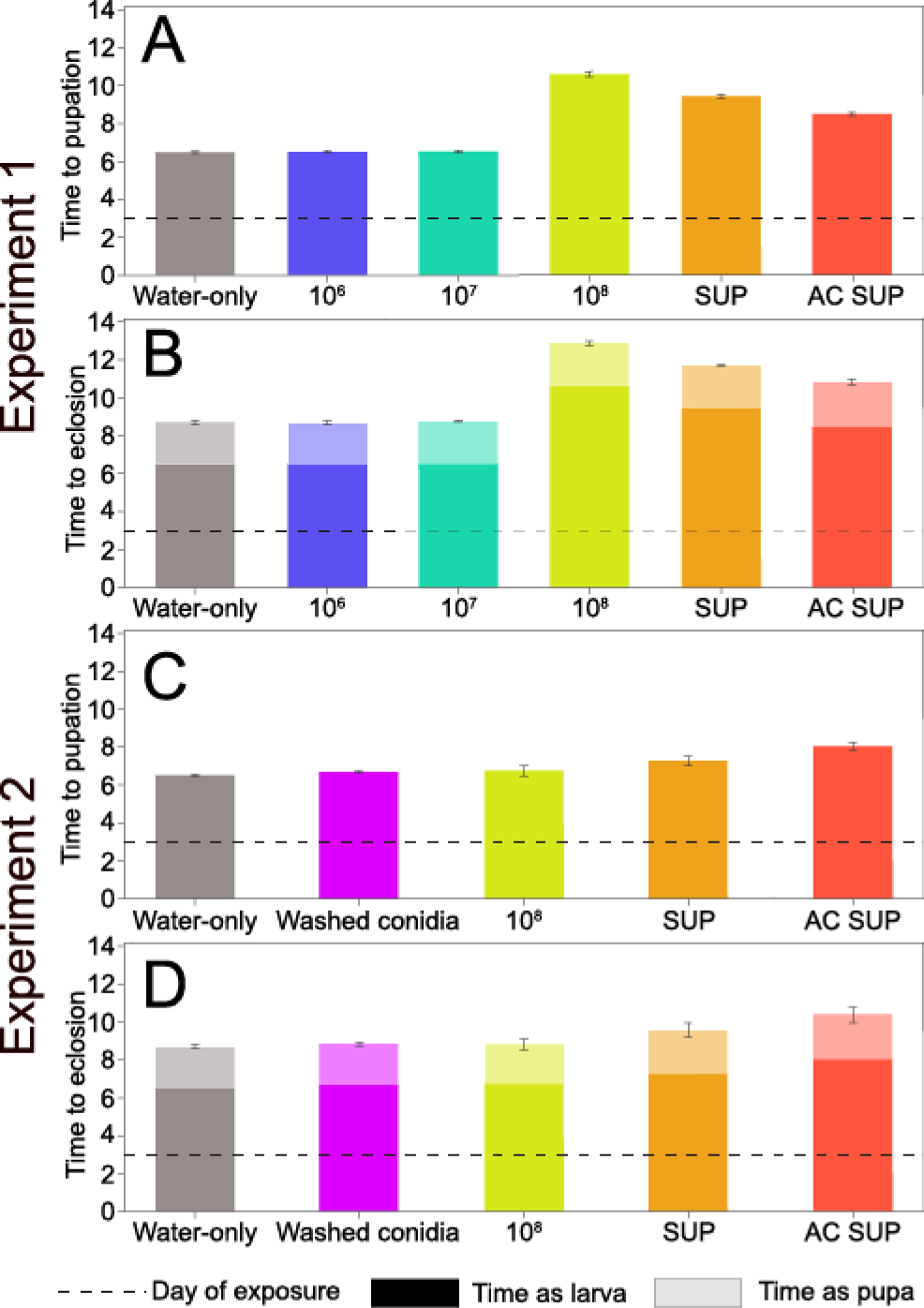
Time to pupation and eclosion of *Ae. albopictus* larvae following exposure to *T. atroviride*. **(A and C)** Mean time to pupation and **(B and D)** eclosion after larval exposure to *T. atroviride* fungal and control treatments. Darker boxes indicate time spent as larva, while lighter color boxes indicate time spent as pupae. Dashed line indicates day of larval exposure. Error bars shown as LSM ± SEM.

**Table 4.**
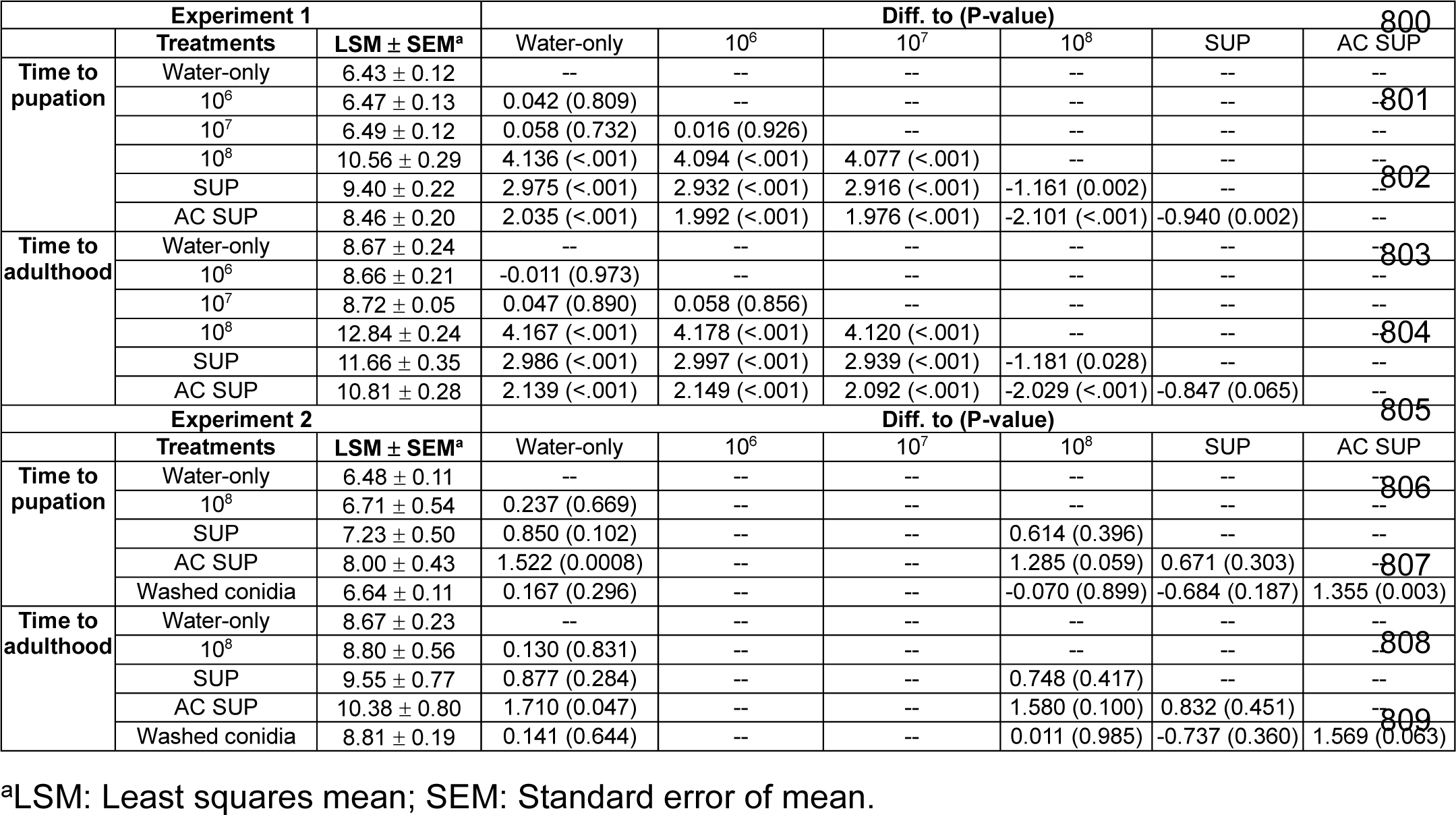
Time to pupation and adulthood of *Ae. albopictus* larvae exposed to *T. atroviride*.

*Aedes albopictus* larvae exposed to the 10^8^ conidia/mL dose and its supernatant treatments developed significantly slower to pupae as compared to the water-only control and lowest conidial doses (Table 4, Fig. 3A). This impact was most strongly observed for larvae exposed to the 10^8^ conidia/mL treatment (4.1 day delay, *P* < 0.001). Significant delays in development time to pupae were also observed with the supernatant (2.97 day delay, *P* < 0.001) and autoclaved supernatant (2.03 day delay, *P* < 0.001) treatments. The delay caused by the 10^8^ conidia/mL treatment and its supernatant was only observed in experiment 1, which likely was the result of the higher larval mortality that we observed in experiment 2 (Tables 1 and 2). While significant delays were observed in time to pupation, adults eclosed 2.1-2.4 days after pupation irrespective of fungal treatment (Table 4, Fig. 3B and D).

### Trichoderma atroviride reduces Ae. albopictus adult fitness

To assess whether larval exposure to *T. atroviride* impacted the fitness of emerging adults, we assessed their survival and body size. To evaluate their survival, we plotted the sex-specific survival proportion of male and female adults ten days post-pupation (Table S3, Fig. 4A and B). Male and female adult survival post-eclosion was not impacted by larval exposure to *T. atroviride* irrespective of treatment. Specifically, male and female adults had survival percentages above 88% across all treatments (Table S3, Fig. 4A and B).

**Fig. 4.**
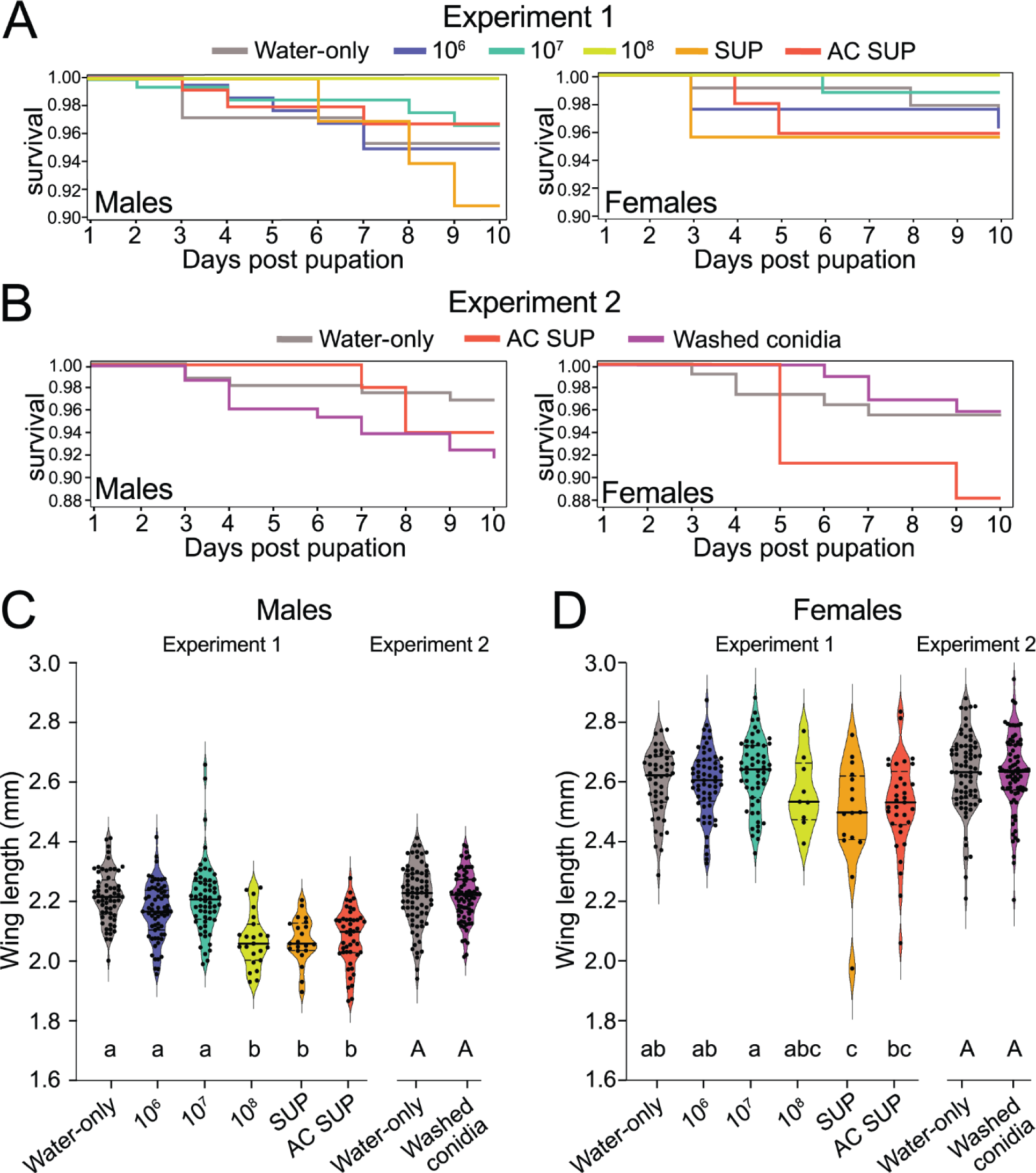
*Aedes albopictus* adult survival and wing length measurements following larval exposure to *T. atroviride*. **(A and B)** Graphs show Kaplan-Meier estimations of survival up to ten days post pupation of adult male and adult females that were exposed as L3 larvae. **(C and D)** Violin plots of adult male and female wing lengths. Males and females were analyzed separately using one-way ANOVA and Tukey’s multiple comparisons test for experiment 1 or t-test for experiment 2. Solid lines show median and dashed lines depict interquartile range.

We then assessed adult wing length as a proxy for adult body size (Fig. 4C and D). We observed significant reduction in the length of male wings when exposed as larvae to the 10^8^ conidia/mL, supernatant, and autoclaved supernatant treatments as compared to the water-only control (F = 20.53; df = 5, 2; *P* < 0.0001, Fig. 4C). The mean wing length of adult males exposed to these treatments was between 2.06-2.09 mm. Males who were exposed to the water-only control as larvae had a significantly higher mean wing length at 2.22 mm. Adult females exposed as larvae to the 10^8^ conidia/mL, supernatant, and autoclaved supernatant treatments had reduced wing lengths compared to the water-only control, though the relative difference was smaller compared to males (F = 4.92; df = 5, 2; *P* = 0.0003, Fig. 4D). Females exposed to the water-only control had a mean wing length of 2.61 mm. A reduction in wing length to 2.49 mm was observed when exposed to the supernatant and 2.53 mm when exposed to the 10^8^ conidia/mL and autoclaved supernatant treatments. We did not observe any difference in the wing length of male or females exposed as larvae to the 10^6^ and 10^7^ conidia/mL or washed conidia treatments.

## Discussion

This study aimed to evaluate the larvicidal potential of a *Trichoderma atroviride* strain isolated from *Aedes albopictus* L4 larvae collected from artificial containers in Manhattan, KS, USA. To do so, we modified a previously established laboratory-based experimental design to evaluate the entomotoxicity of *T. atroviride* culture supernatant and entomopathogenicity of *T. atroviride* conidia (washed conidia treatment) on *Ae. albopictus* larvae (Tawidian et al., 2023). While the efficacy of fungal bioinsecticides in the field are impacted by a variety of abiotic and biotic factors (Fang and St. Leger, 2012; Sharma et al., 2020), this study establishes that *T. atroviride* culture supernatant is a potent entomotoxin towards *Ae. albopictus* larvae.

This study did not find any evidence of entomopathogenicity of *T. atroviride* conidia against *Ae. albopictus* larvae. The culture supernatant without conidia killed larvae to the same level as the culture supernatant with conidia and conidia alone did not kill larvae, despite conidia remaining viable throughout the exposure period and being ingested. We cannot exclude that lack of infection may be the result of *T. atroviride* conidia not being able to interact with larval cuticle over expanded periods of time, as conidia are hydrophobic, tend to clump together and thus sink in the water column (Bukhari et al., 2010; Cai et al., 2020). However, the mosquito larval cuticle and gut epithelium are themselves challenging barriers that conidia must adhere to and penetrate rapidly, as mosquito larvae purge their gut content before molting and fungi may be shed with the exuvium during molting (Vestergaard et al., 1995). This is especially true for *Ae. albopictus,* as they proceed rapidly through their larval development. Regardless of the underlying reasons for the lack of observed entomopathogenicity, we show definitively that the observed larvicidal activity of *T. atroviride* is due to entomotoxicity of its culture supernatant.

*Trichoderma atroviride* culture supernatant affected mosquito fitness by killing mosquito larvae. High concentration fungal treatments, containing water-soluble metabolites, significantly reduced larval survival to the pupal and adult stages while disproportionately targeting females. Our results parallel studies that report the larvicidal impact of culture supernatant from other *Trichoderma* species towards several mosquito species, including *Ae. albopictus*, *Aedes aegypti*, *Culex quinquefasciatus*, and *Anopheles* species (Govindarajan et al., 2005; Rana and Sandhu, 2008; Podder and Ghosh, 2019; da Silveira et al., 2021; Perera et al., 2023). To our knowledge, this is the first report of the larvicidal potency of the culture supernatant of a *T. atroviride* strain towards mosquitoes. Up to this point, the only species for which *T. atroviride* metabolites were shown to exhibit entomotoxicity was *Drosophila melanogaster* (Atriztán-Hernández et al., 2019). A previous study in *Ae. aegypti* showed that silver nanoparticles incubated with *T. atroviride* culture filtrate were toxic to larvae, but did not examine whether toxicity was conferred by culture filtrate alone (Singh and Prakash, 2015). Our observation that female larvae are killed at a higher proportion than male larvae is likely a consequence of their larger size (Armbruster and Conn, 2006), which we also observed in our experimental setup. This not only provides more larval surface area of exposure to contact toxins, but also requires female larvae to ingest more food, including higher amounts of both conidia and culture supernatant.

In addition to killing, *T. atroviride* culture supernatant further reduced mosquito fitness by extending larval development time. Similarly, exposure to *B. bassiana* and *M. anisopliae* conidia have been shown previously to delay larval development of *Anopheles stephensi* and *Culex pipiens* (Hamama et al., 2022; Renuka et al., 2023). The developmental delay we observe resembles exposure of mosquito larvae to sublethal doses of chemical or biological larvicides. For example, time to pupation for *Ae. aegypti* larvae exposed to *B. thuringiensis* increased by more than four days compared to their untreated counterparts (Wang and Jaal, 2005). Several metabolites synthesized by *Trichoderma* species, including volatile organic compounds and chitinases, have delayed larval development in *D. melanogaster* and *Bombyx mori*, respectively (Inamdar et al., 2012; Berini et al., 2015). It is possible that these metabolites are present in the culture supernatants used in this study and exert sublethal effects on *Ae. albopictus* (e.g. midgut epithelial damage) that delays larval development.

In addition to larval killing and extension of development time, *T. atroviride* culture supernatant further reduced mosquito fitness by reducing adult size. While adult survival was not impacted, both male and female wing lengths following exposure to culture supernatant were reduced significantly. Adult wing length is a standard proxy for adult body weight in mosquitoes, which is positively correlated with adult mosquito fitness in both sexes. For example, adult size effects sperm capacity (Ponlawat and Harrington, 2007; Helinski and Harrington, 2011) and male mating success (Yuval et al., 1993; Sawadogo et al., 2013), as well as blood meal size, biting frequency (Scott et al., 2000; Phasomkusolsil et al., 2015), and egg count (Briegel and Timmermann, 2001; Armbruster and Hutchinson, 2002; Farjana and Tuno, 2012; Yamany and Abdel-Gaber, 2024). Based on this literature, the observed reduction in wing length of *Ae. albopictus* may have pleiotropic effects on adult mosquito fitness potentially affecting feeding behavior, mating success, and fecundity. Future studies will have to decipher which if any of these *Ae. albopictus* adult fitness parameters are affected by larval exposure to *T. atroviride*.

We found that the observed entomotoxicity of *T. atroviride* against *Ae. albopictus* is due to a mixture of heat-stable and heat-labile metabolites found in the culture supernatant. Larval killing, delayed development time, and adult body size were reduced significantly when culture supernatants were heat-treated by autoclaving, indicating that heat-stable entomotoxins contribute to the observed effects. *Trichoderma* species produce many toxins with insecticidal or antifeedant properties, such as volatile organic compounds (e.g., α-pyrones), peptaibols, and digestive enzymes (e.g., chitinases and proteases) (Poveda, 2021). Notably, peptaibols have high thermal stability (Song et al., 2006; De Zotti et al., 2020) and therefore likely contribute to the observed larvicidal activity of the autoclaved *T. atroviride* culture supernatant.

In conclusion, we report the identification of a mosquito-associated *T. atroviride* strain with potential for *Ae. albopictus* larval control. Future studies should evaluate existing commercial formulations of *T. atroviride* used in agricultural settings for their efficacy against mosquito larvae and identify the entomotoxins present in the culture supernatant that exhibit larvicidal activity.

## Supporting information

Supplementary Information TablesS2,3 & Figures S1-5

Supplementary Table S1: Raw data

## Acknowledgements

We acknowledge that larval collections and our research were conducted in Manhattan, KS, on land historically home to many Native nations, including the Kaw, Osage, and Pawnee, among others. We further acknowledge that Kansas State’s history as the first land grant university in the United States rests on the dispossession of Indigenous peoples and nations from their lands. We thank Jordan Block, Kansas State University, for technical help with mosquito rearing and fungal exposures.

## Author Contribution Statements

DTH: Conceptualization, Data curation, Formal analysis, Investigation, Methodology, Software, Validation, Visualization, Writing – original draft, Writing – review & editing; PT: Conceptualization, Data curation, Investigation, Methodology, Resources, Validation, Visualization, Writing – original draft, Writing— review & editing; ES: Formal analysis, Methodology, Software, Validation, Writing— review & editing. QK: Methodology, Software Writing— review & editing; AJS: Investigation, Writing – review & editing; KM: Conceptualization, Data curation, Funding acquisition, Methodology, Project administration, Resources, Supervision, Validation, Visualization, Writing – original draft, Writing – review & editing.

## Funding

KM and QK are members of the Vector-borne and Parasitic Disease team at Kansas State, which was supported by an internal Game-changing Research Initiation Program seed grant from the Office of the Vice President of Research. This study was supported by funding to KM from National Institute of Allergy and Infectious Disease grant R01AI140760, Agricultural Research Service specific cooperative agreement 58-5430-4-022, and National Institute of Food and Agriculture Hatch projects 1021223 and 7007965. This is contribution 25-134-J from the Kansas Agricultural Experiment Station. The contents of this article are solely the responsibility of the authors and do not necessarily represent the official views of the funding agencies.

## Notes

### Competing Interest Statement

The authors have declared no competing interest.

