## Supplementary Information TablesS2,3 & Figures S1-5 for "Larvicidal activity of *Trichoderma atroviride* (Hypocreales: Hypocreaceae) against *Aedes albopictus* (Diptera: Culicidae)"

**Supplementary Materials:**

**Supplementary Tables S2-3** (Please note that Supplementary Table S1 is provided as an Excel file only)

**Supplementary Figures S1-5**

**Table S2. Survival of *Ae. albopictus* pupae to the adult stage.**

| Experiment 1 | Treatments | Survived to adulthood | Died as pupa | Percent mortality | *P* |
|---|---|---|---|---|---|
| | Water-only | 191 | 13 | 6.373 | -- |
| | $10^6$ | 188 | 14 | 6.931 | 0.845 |
| | $10^7$ | 193 | 9 | 4.455 | 0.512 |
| | $10^8$ | 54 | 9 | 14.286 | 0.064 |
| | SUP | 49 | 9 | 15.517 | 0.034 |
| | AC SUP | 133 | 20 | 13.072 | 0.041 |
| **Experiment 2** | **Treatments** | **Survived to adulthood** | **Died as pupa** | **Percent mortality** | ***P*** |
| | Water-only | 269 | 10 | 3.584 | -- |
| | $10^8$ | 6 | 1 | 14.286 | -- |
| | SUP | 19 | 2 | 9.524 | 0.202 |
| | AC SUP | 91 | 9 | 9.000 | 0.057 |
| | Washed conidia | 264 | 15 | 5.376 | 0.414 |

**Table S3. Hazard ratio of male and female *Ae. albopictus* adults up to ten days post-pupation following surviving larval exposure to *T. atroviride*.**

| Experiment 1 | | Hazard Ratio. to (P-value) | | | | |
|---|---|---|---|---|---|---|
| | **Treatments[a]** | Water-only | $10^6$ | $10^7$ | SUP | AC SUP |
| **Female** | Water-only | -- | -- | -- | -- | -- |
| | $10^6$ | 1.360 (0.689) | -- | -- | -- | -- |
| | $10^7$ | 0.317 (0.321) | 0.233 (0.193) | -- | -- | -- |
| | SUP | 1.534 (0.712) | 1.128 (0.915) | 4.835 (0.269) | -- | -- |
| | AC SUP | 0.989 (0.990) | 0.727 (0.716) | 3.115 (0.358) | 1.552 (0.720) | -- |
| **Male** | Water-only | -- | -- | -- | -- | -- |
| | $10^6$ | 1.003 (0.996) | -- | -- | -- | -- |
| | $10^7$ | 0.625 (0.467) | 0.623 (0.464) | -- | -- | -- |
| | SUP | 1.454 (0.598) | 1.449 (0.602) | 2.326 (0.270) | -- | -- |
| | AC SUP | 0.539 (0.383) | 0.537 (0.383) | 0.862 (0.847) | 2.697 (0.225) | -- |
| **Experiment 2** | | **Hazard Ratio. to (P-value)** | | | | |
| | **Treatments[b]** | Water-only | $10^6$ | $10^7$ | SUP | AC SUP |
| **Female** | Water-only | -- | -- | -- | -- | -- |
| | AC SUP | 0.585 (0.442) | -- | -- | -- | -- |
| | Washed conidia | 1.629 (0.473) | -- | -- | -- | 2.786 (0.168) |
| **Male** | Water-only | -- | -- | -- | -- | -- |
| | AC SUP | 0.579 (0.476) | -- | -- | -- | -- |
| | Washed conidia | 0.355 (0.054) | -- | -- | -- | 0.614 (0.473) |

[a]$10^8$ conidia/mL treatment excluded from hazard analyses as no adult death was observed. [b]$10^8$ conidia/mL and supernatant treatments excluded from hazard analyses as too few adults emerged to evaluate.

**Supplementary figures with legends**

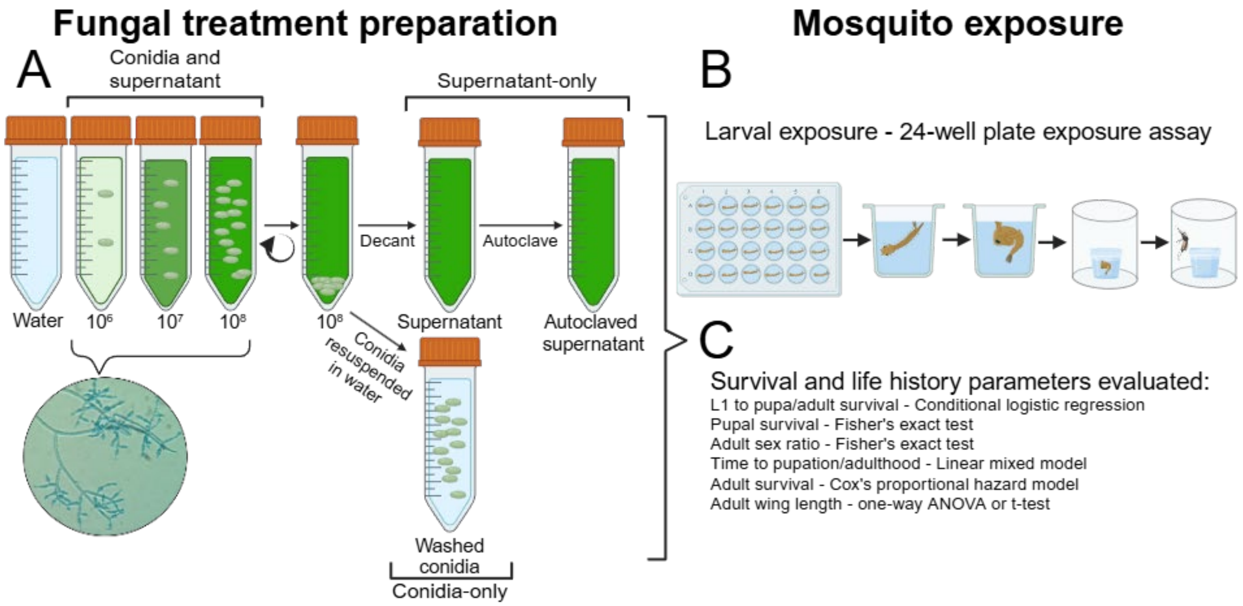


**Fig. S1. Graphical methods of *Trichoderma atroviride* treatment preparations and larval exposure. (A)** Fungal conidia and culture supernatant were collected from wheat kernels to prepare treatments. The $10^6$ and $10^7$ conidia/mL treatments were included only in experiment 1, while the washed conidia treatment was included only in experiment 2. **(B)** Larval exposure of fungal and control treatments. **(C)** Survival and life history parameters evaluated and their respective statistical analyses. Created in https://BioRender.com

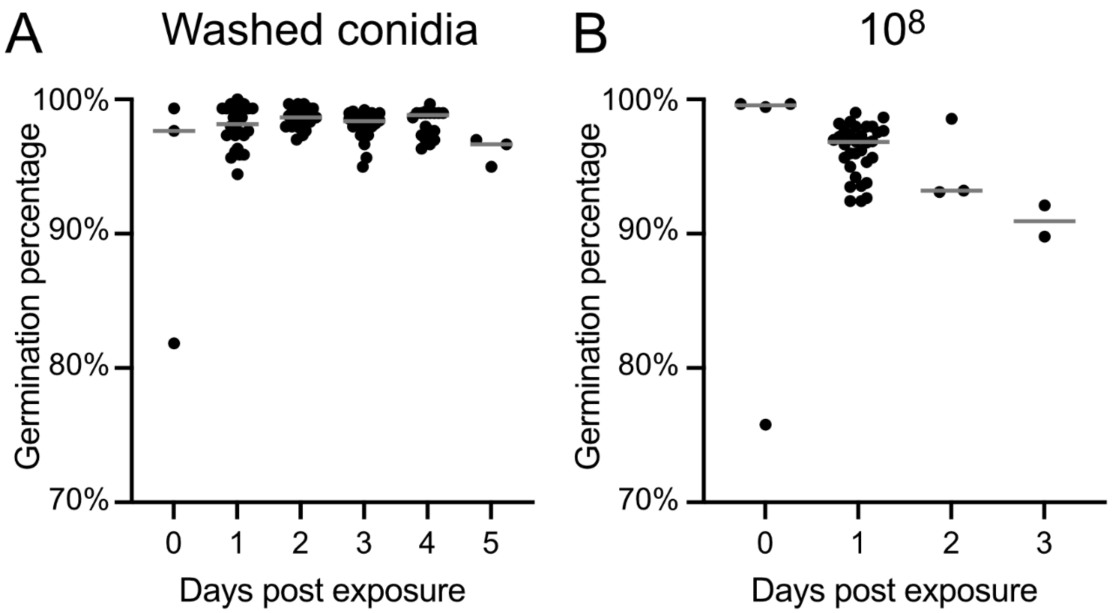


**Fig. S2. Daily germination percentage of *T. atroviride* conidia in larval microcosms**. **(A)** Washed conidia treatment, **(B)** $10^8$ conidia/mL treatment. Larvae were exposed to $10^8$ conidia on day 0. Differences in germination percentages over time for washed conidia were assessed using a linear regression model with Days post exposure as predictor variable and Germination percentage as response variable (df = 94, Residual std. error = 0.002, Adj $R^2$ = 0.0023, F = 1.216, $P$ = 0.2729). Median line shown in gray.

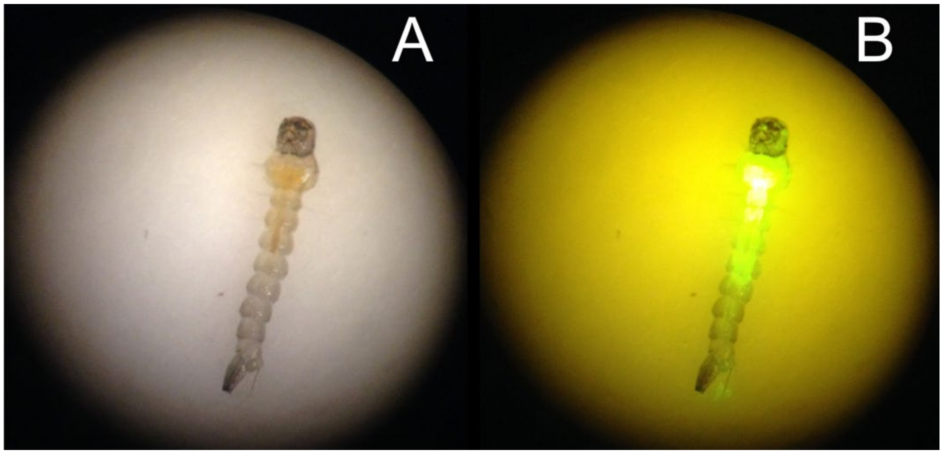


**Fig. S3. *Aedes albopictus* larvae consumed *T. atroviride* conidia.** *Aedes albopictus* larvae fed for 24 hours with acridine orange-labelled *T. atroviride* washed conidia. **(A)** Image under brightfield, **(B)** Image under eGFP filter.

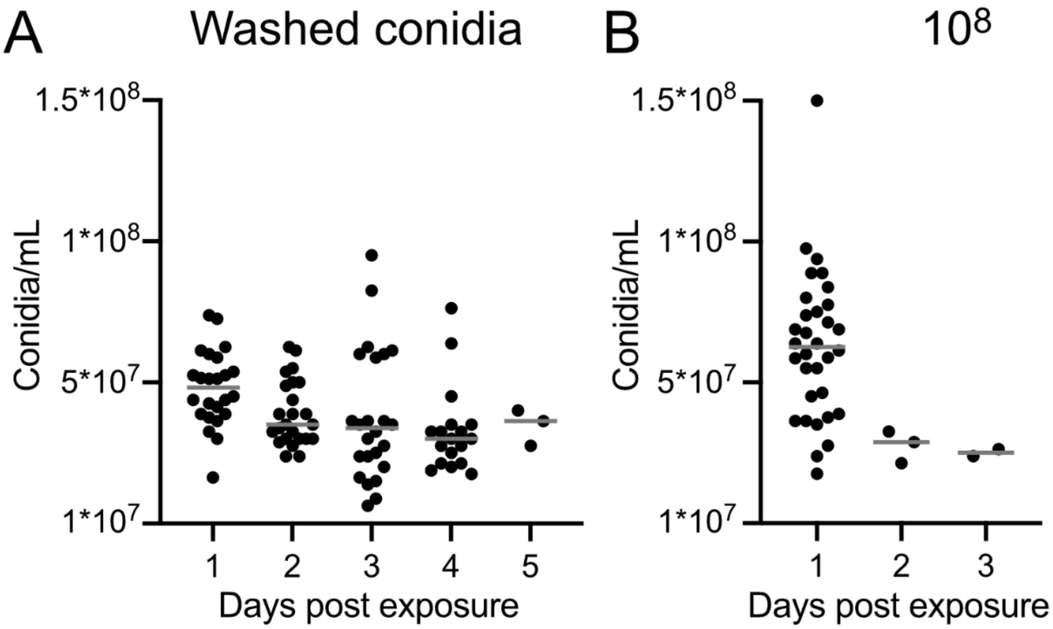


**Fig. S4. Daily conidia concentration of *T. atroviride* in larval microcosms**. **(A)** Washed conidia treatment, **(B)** $10^8$ conidia/mL treatment. Larvae were exposed to $10^8$ conidia on day 0. Differences in conidial concentrations over time were assessed using a linear regression model with Days post exposure as predictor variable and Conidia/mL as response variable (df = 91, Residual std. error = $1.623x10^7$, Adj R2 = 0.0762, F = 8.548, $P$ = 0.0043). Median line shown in gray.

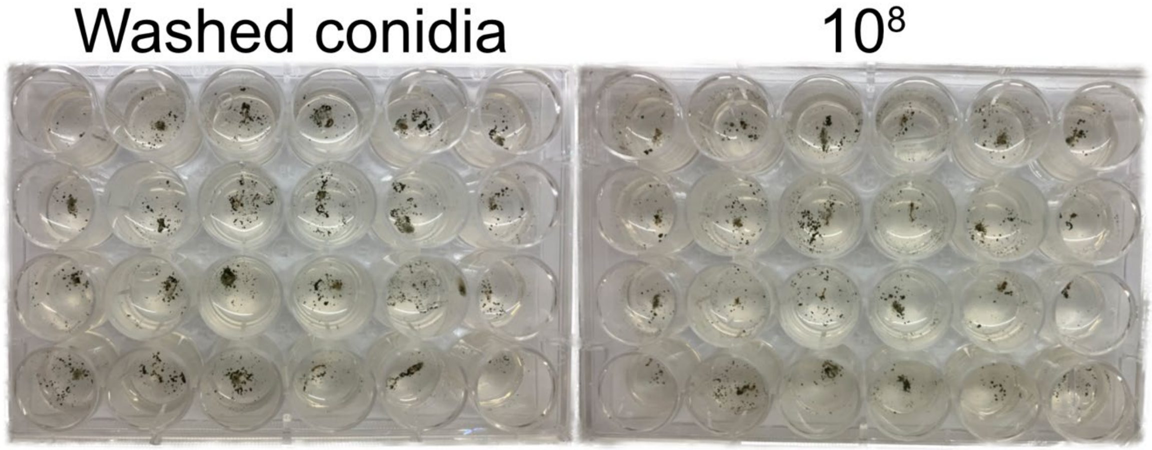


**Fig. S5. Sporulation of *T. atroviride* conidia from *Ae. albopictus* larval carcasses.** Dead larvae exposed to washed conidia and $10^8$ conidia/mL treatments were surface-sterilized and individually plated into wells of a 24 well plate containing agar lacking any nutrients.
